# Administration of defective virus inhibits dengue transmission into mosquitoes

**DOI:** 10.1101/837849

**Authors:** Tarunendu Mapder, John Aaskov, Kevin Burrage

**Affiliations:** School of Mathematical Sciences, Queensland University of Technology, Brisbane, Queensland, Australia; Australian Research Council Centre of Excellence for Mathematical and Statistical Frontiers, Queensland University of Technology, Brisbane, Queensland, Australia; Institute of Health and Biomedical Innovation, Queensland University of Technology, Brisbane, Queensland, Australia; Department of Computer Science, University of Oxford, Oxford, UK

## Abstract

The host-vector shuttle and the bottleneck in dengue transmission is a significant aspect with regard to the study of dengue outbreaks. As mosquitoes require 100-1000 times more virus to become infected than human, the transmission of dengue virus from human to mosquito is a vulnerability that can be targeted to improve disease control. In order to capture the heterogeneity in the infectiousness of an infected patient population towards the mosquito pool, we calibrate a population of host-to-vector virus transmission models based on an experimentally quantified infected fraction of a mosquito population. Once the population of models is well-calibrated, we deploy a population of controls that helps to inhibit the human-to-mosquito transmission of the dengue virus indirectly by reducing the viral load in the patient body fluid. We use an optimal bang-bang control on the administration of the defective virus (transmissible interfering particles, known as TIPs) to symptomatic patients in the course of their febrile period and observe the dynamics in successful reduction of dengue spread into mosquitoes.

## Introduction

Dengue is a major health burden in tropical regions ^1, 2^, which is caused by the systematic and self-limited transmission of dengue virus (DENV) between human and mosquitoes ^3^. Four serotypes of the DENV arbovirus have been identified as being transmitted by *Aedis aegypti* mosquitoes ^4^. Although four serotypes have different interactions with the host antibodies, they result the same disease with the same clinical symptoms. As *Aedis aegypti* is a competent vector for all dengue serotypes, DENV can systematically infect and persist within the mosquito body, and eventually the vector becomes infectious ^5–8^. The presence of DENV in the abdomen or saliva defines a mosquito to be infected or infectious, respectively. However, the passage of DENV from the mosquito mid-gut to the salivary gland needs to cross a number of tissues barriers and innate immunity responses. To cross these hurdles, DENV needs to replicate across all the different tissues and body fluids ^9, 10^. For this reason, a bottleneck arises during the transmission flow from the human host to the mosquitoes. We suggest this phenomenon as a potentially efficient dengue control.

Traditional vector controls based on reducing the mosquito population have not been successful in endemic and pandemic situations for dengue ^11^. Destroying mosquito habitats using insecticides and biological controls by nematodes and fungi are primitive approaches for vector control. Due to the lack of success of these approaches ^12, 13^, genetic manipulation and population replacement of mosquitoes has recently been attempted ^3, 14, 15^. This has not been successful due to the inability of these populations to invade the wild mosquito population. In consequence, the global spread of dengue is pandemic, accentuated, for example, through increased travel between countries and urban growth ^16, 17^.

Another popular control of infectious diseases spread is through vaccination. Intervention with an attenuated live virus has been successfully implemented in eradication of some diseases such as smallpox ^18^. Although vaccines are available for infectious diseases including dengue, the approach has limited success due to limited vaccine coverage, obstruction in herd immunity and financial burden ^19–21^. However, the most crucial reason of the failure of the conventional vaccination programs for RNA viral diseases is the frequent mutation and fast evolution rate. Thus, an efficient vaccination strategy should be co-evolving and persist over passages and through the host-vector transmission shuttle.

Defective interfering (DI) viruses, containing neucleotide deletions, can be deployed as a potential co-evolving intervention. In the case of RNA viruses, defective viral genomes (DVG) are spontaneously generated within infected host cells ^22, 23^. Upon infection, the positive-strand RNA appears as mRNA for translation of the replication and encapsidation machinery. The dynamic secondary and tertiary structures of positive-strand RNA plays a crucial role in regulation of translation, transcription, replication and encapsidation ^24^. Because of the RNA structure-mediated binding of the polymerase, ‘snapback’ and ‘copyback’ replication mechanisms can generate defective genomes ^25^. Within infected host cells, the defective RNAs interfere with virus replication and help attenuate the plasma viraemia level in the host body ^26, 27^. The DI particles are so named because of the interference with the multiplication of standard virus particles. The DI particles and standard virus stimulate the host immune responses in the same way as the DI particles are encapsidated by the normal coat proteins synthesised by the standard virus ^28^. There is a large number of models studied on the generation and activities of DI particles for many animal viruses ^29, 30^.

We have recently proposed a within-host model of dengue virus infection that covers the natural synthesis of dengue DI particles ^31^. We have constructed a population of models (POMs), calibrated with clinical data for 208 patients ^32^. The intention of the POMs study is to capture the biological heterogeneity in, for example, patient response, which is difficult to investigate with experimental methods alone ^33^. One cause of variability is the cellular infection responsible for the viral load in the patients’ blood samples. Infection and super-infection by standard virus and DI particles introduce variability at numerous stages, including the membrane receptor dynamics of early-infected cells, and activation of the interferon pathways in the late-infected cells ^34, 35^. Four clinically calibrated serotype-specific POMs have been employed to investigate variability in dengue infection in physiological and pathological conditions. In the same framework, we have discussed the effect of adding excess DI particles through a bang-bang optimal control as an intervention strategy. Addition of excess DI particles to a host system that naturally generates DI particles, helps to reduce plasma viraemia peak and duration. Hence, we have predicted four serotype-specific populations of DI particles-mediated controls. We identify the population of vectors of doses and durations of DI particles addition as population of controls (POCs). Naturally, we observe attenuation of the viraemia peaks and duration after successful deployment of the POCs and now the POMs are known as the controlled POMs (cPOMs). The POCs have also been characterised on the basis of its efficiency in reduction of the within-host viral burdens.

As discussed before, the transmission of dengue virus in mosquitoes is highly sensitive to the viral load of the infected patients. In this paper, we elucidate the dynamics of dengue virus transmission from infected patients into a mosquito population. Hence, we demonstrate the fraction of viral infected mosquito population shaped by the host viraemia profile. The timeframe of successful transmission is described at a demographic scale. We hypothesize that controlling the plasma viraemia within the infected host may efficiently restrict the host-to-vector transmission and hence the outbreak of dengue.

## Methods

The key aim of this paper is to study the transmission dynamics of DENV from human host to vector (mosquito) and the effect of adding excess DI particles on the virus transmission. To validate and calibrate the models, we utilize a set of clinical data for 208 adult dengue patients, collected by Nguyen *et. al*. in Vietnam ^32^. They recorded the plasma serology profiles and quantified viraemia levels on a daily basis. In a previous study, we have deployed the patients’ blood sample data to calibrate a population of within-host models (POMs) and predicted a population of personalised controls (POCs) that is able to reduce blood viraemia levels ^31^. Nguyen *et. al*. also performed a mosquito exposure experiment on the same group of patients. With informed consent, each patient has been assigned for multiple exposures to mature and competent mosquitoes. 408 exposures were scheduled purely randomly during high fever days. In each exposure, approximately 25-40 female 3-7 days old *Aedis aegypti* mosquitoes were allowed to bite on the patient’s forearm. We use this dataset of mosquito exposures to help us explore the host-to-vector transmission dynamics. In this paper, we model the mosquito exposure experiment to occur in an open environment within the territory of a mosquito pool. Hence, an experimentally calibrated population of transmission models is proposed for each dengue serotype. Here we note that the serotype-specific POMs are calibrated only based on the recorded dataset, otherwise the models follow the same mathematical structure. We are unable to incorporate any serotype-specific interaction based on the available data of viral load in the mosquito abdomen and patients’ serology.

### Within-host Dengue viraemia

The data of virus and DI particles in the patients’ blood samples are used as input for the present model. The kinetics of plasma viraemia and their defective counterpart are estimated from a within-host dengue virus infection model. The within-host model describes the dynamics of cell-virus interaction and the triggered immune response inside the human host body. As the DI particles can replicate only in the dually infected cells, these cells are the only source in which to accumulate DI particles in the host body fluid. Hence, every time an infected host with high plasma viraemia is exposed to mosquitoes, there is a possibility that the mosquitoes while taking a blood meal are infected in three different ways: by the virus, by the DI particles or dually. As we have also predicted successful optimal bang-bang control strategy for patients’ viraemia reduction, the controlled viraemia profiles are available from the within-host model ^31^. The controlled profiles are used in the present model to observe their efficiency in transmission reduction.

### Dengue transmission in mosquitoes

With the natural birth-death flow of mosquitoes this model examines how a population of mosquitoes take blood meal from infected patients on different days of their febrile period in a continuous manner. The reported data were collected on specific days, on which the mosquitoes are harvested for quantitative assays. Hence, we found a daily distribution of infected mosquitoes in order to calibrate our population of models. In general, many mathematical models of infectious disease spread follow the very well-established SIR model ^36, 37^. Variants include adding compartmental models, seasonal effects and spatio-temporal dynamics ^37–39^. However, tracking the transmission of the host plasma viral load to the mosquito pool within the febrile period has not yet been explored. We follow the traditional SIR approach but do not include the recovered (R) compartment, as mosquitoes die naturally before they can recover from dengue virus infection. We construct a population of transmission model that utilizes the patient viremia and DI particles profiles from our previous model ^31^. This model can explain the infected fraction of the mosquito population in terms of the transmission of viremia from the infectious patients. The mathematical structure of the model is shown below.

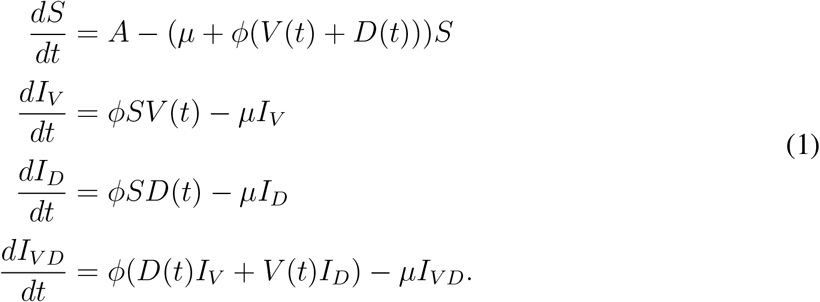

Here, *S*, *I*_*V*_, *I*_*D*_, and *I*_*V D*_ are the susceptible and three kinds of infected pools of mature mosquitoes, respectively. The infected pools are by virus only (*I*_*V*_), by DIPs only (*I*_*D*_) and by both virus and DI particles (*I*_*V D*_). *A*, *ϕ* and *μ* are the rate of reproduction, infection and natural death of mosquitoes, respectively. The plasma viral load (*V*(*t*)) and DI particles (*D*(*t*)) are obtained from our previous study ^31^.

### Population of models

Characterising the variability between individuals from the same species is fundamental in biology. In our setting, every infected individual in a population may not produce similar infectiousness to the similar groups of susceptible, mature mosquitoes. The underlying variability may be manifested by: (1) the patients’ physiology and immunological response, (2) behaviour of the uptaken virus on interaction with the host cells, (3) mosquito physiology. The first two points are covered when we use a calibrated population of patients’ viral load (*V*(*t*)) and associated DI particles (*D*(*t*)). To capture the variability in patient infectiousness towards mosquitoes, a population of models approach is used for the transmission model. The main purpose of such a study is to decipher how a particular population of patients are different from the others (i.e., infectiousness to mosquitoes at different levels of viral load) and on the clinical realm, compared with the treated (controlled) population. In a clinical and experimental framework, it is difficult to track the variability, so population of mathematical models can overcome this problem even in low sample size. Two patients with similar viraemia profiles, for example, when exposed to the same mosquito pools may show very different infectiousness. In such cases, the use of a variable population of models is more realistic than a single mean-field model ^40, 41^.

The population of transmission models takes the form in Eq. 1 and is calibrated by sampling the model parameters based on Latin Hypercube Sampling (LHS) ^42^. Uniformly selected 15% profiles of viremia (*V*(*t*)) and DI particles (*D*(*t*)) from the infected patients’ population are used here as model input and the parameters (*A*, *μ* and *ϕ*) are sampled from Latin Hypercube Sampling. We are free to set different criteria for our model calibration. A previously reported article has chosen the range of the biomarker data as the calibration criteria ^40^, which is sometimes coarse. In our recent article we have calibrated models by matching the distribution of the data ^43^. As distribution matching is computationally expensive and an initial observation did not offer significant insight in our present study, we adopt the previous range-based method as described in Algorithm 1. The model calibration process is based on the range of the experimental biomarker data ^31, 33^. The models, which produce the daily infected mosquito pool within the range of reported data, are included into the population from an initial population of 1,000 sampled models for each individual patient model. We construct four separate populations of the different fractions of infected mosquitoes pool for four serotypes of DENV.

**Algorithm 1.**
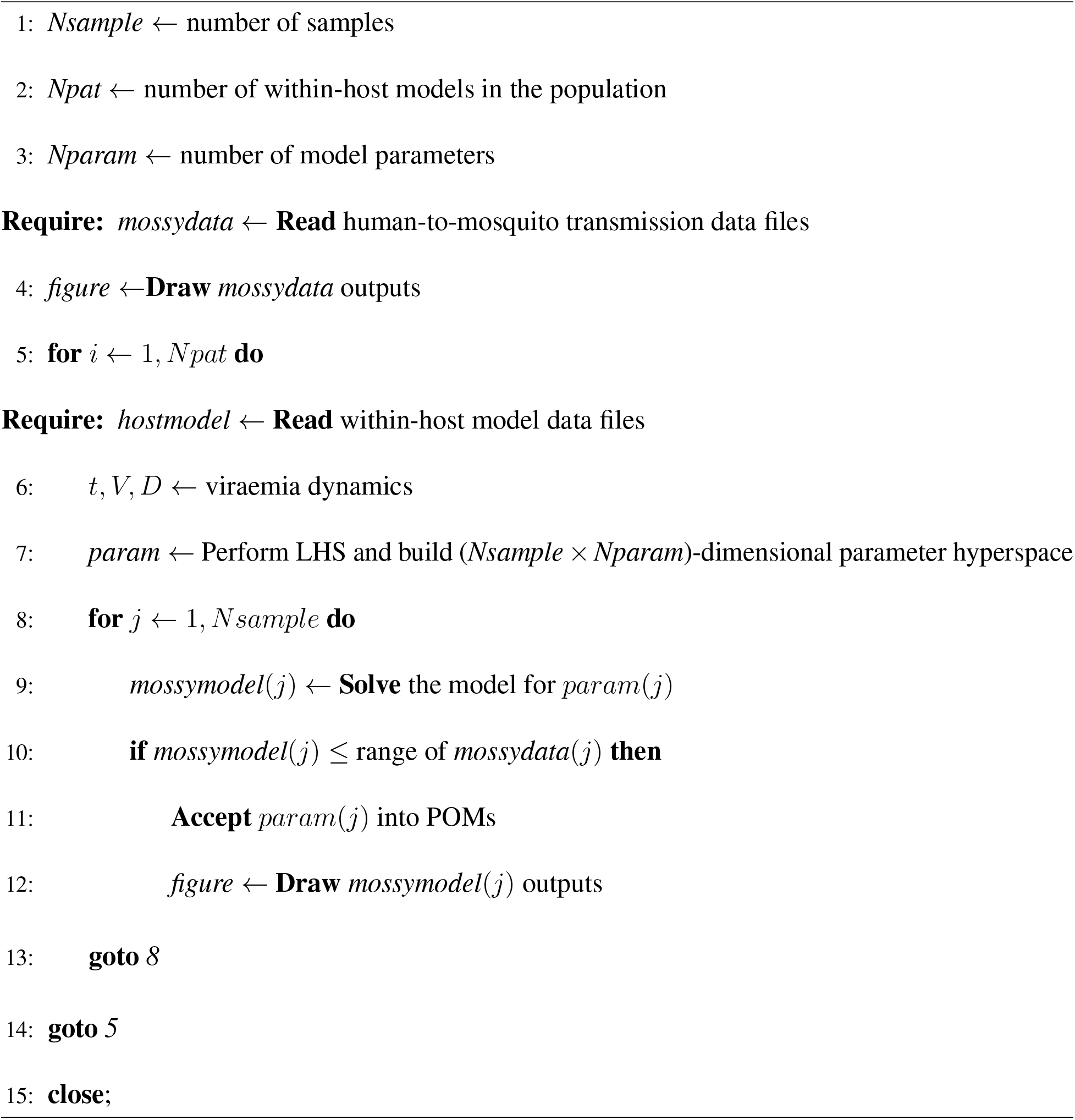
Construction of experimentally calibrated transmission POMs

### Optimal bang-bang control

In the present study, we use the output from the same population of within-host models twice; first, the uncontrolled *V*(*t*) and *D*(*t*) while constructing the infection population of mosquitoes and secondly, the controlled *V*(*t*) and *D*(*t*), which leads to high reduction of the host-vector transmission of dengue virus. Here the controlled *V*(*t*) and *D*(*t*) appear after application of an optimal bang-bang control in the within-host POMs ^31^. The purpose of optimal control is to determine the optimal trajectory of a control variable over time by optimising a predefined objective function using dynamical programming. The Pontryagin’s minimum principle and Hamilton-Jacobi-Bellman theorem are popularly used to solve dynamical control problems ^44, 45^. In dynamical programming, optimal controls are of two kinds: continuous and bang-bang. Although continuous optimal control is popular in engineering and biology, bang-bang control is less popular due to computational difficulties. Bang-bang control flips between the ‘on’ and ‘off’ states depending on the states of the system; hence it is more relevant in clinical interventions. Recently, we have discussed the successful implementation of both continuous and bang-bang optimal control to predict interventions in disease models such as acute myeloid leukaemia and dengue infection ^31, 46^.

## Results

The infected mosquitoes are classified according to the nature of infecting agents as: infected by virus (*V*(*t*)), infected by DI particles (*D*(*t*)) and infected by both *V*(*t*) and *D*(*t*). We denote each fraction of the infected mosquitoes with respect to the entire mosquito pool as

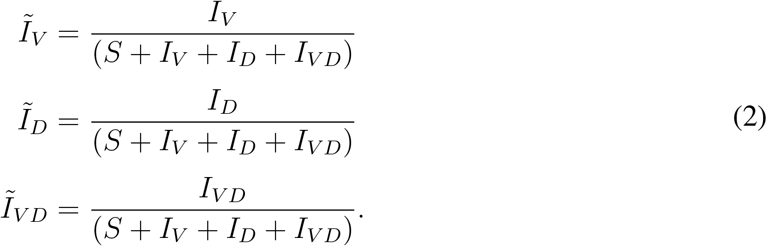

The profiles of 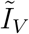, 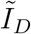 and 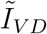 are shown in Fig 2. The 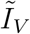 fraction is calibrated using the available clinical data, shown in black dots for 208 patients, assigned in 408 mosquito exposure experiments. On an average, every patient has been exposed twice within day 2 and day 7. The other two panels, 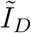 and 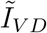, are estimated from the simulated models while calibrating the POMs. For each input patient’s model with a particular serotype, we simulate the same model (Eq 1) for 1,000 sampled sets of parameters from Latin Hypercube Sampling and select only those models that appear with 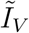 levels within the ranges of the data. The allowed ranges of the parameters are: *A* → (5.0 × 10^−3^ − 50.0) per day, *μ* → (1.567 × 10^−4^ – 1.567) per day and *ϕ* → (7.5 × 10^−11^ – 7.5 × 10^−7^) per day, which are within the biologically feasible ranges ^37^. The patient-specific POMs are shown in different colours in Fig 2. These calibrated populations of transmission models are simulated again with the treated (controlled) viraemias. The treated viraemias are obtained from the corresponding within-host models after applying bang-bang optimal control. As the controlled within-host viraemia are highly reduced, the transmission of the viraemia into mosquito pool are also significantly low (see Fig 5).

**Figure 1:**
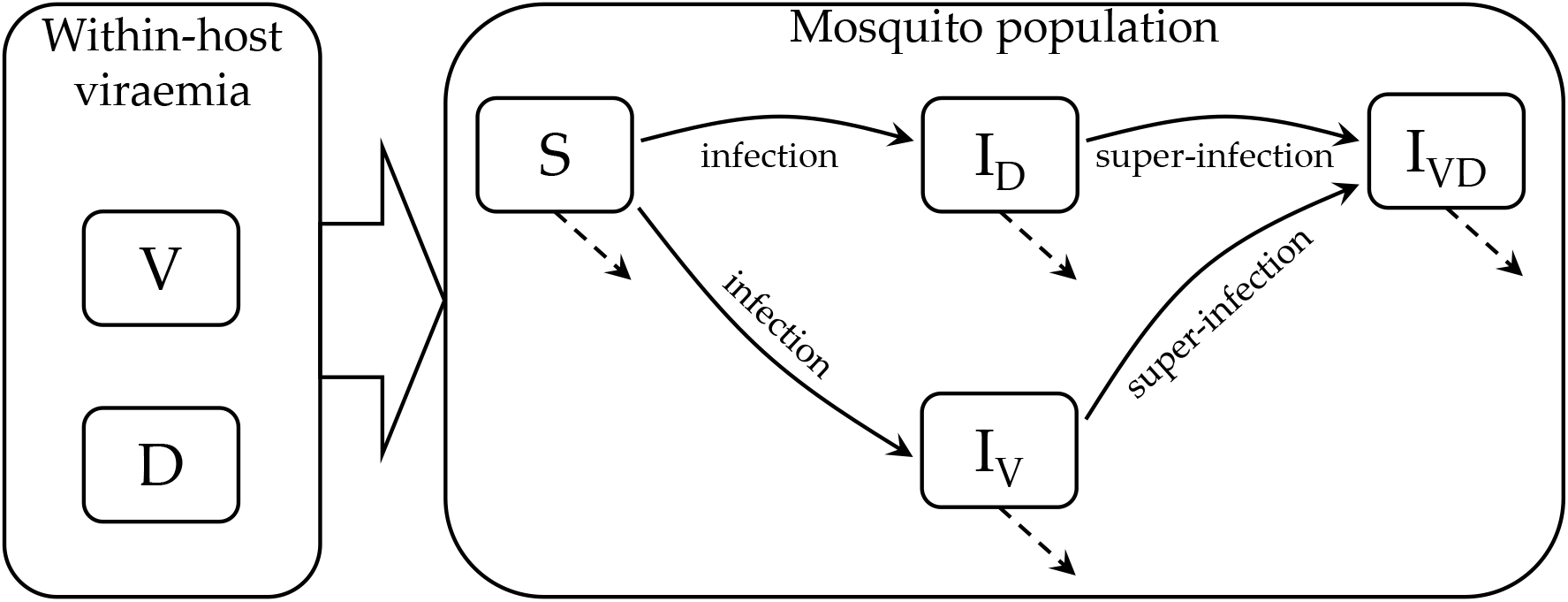
Schematic diagram of dengue virus transmission model: the virus (*V*) and the DI particles (*D*) are transmitted from infected hosts to mosquitoes. The susceptible mosquitoes (*S*) can be infected by virus or DI particles to generate the two types of infected mosquitoes, *I*_*V*_ and *I*_*D*_. Further, a super-infected pool (*I*_*V D*_) is generated from further infection of *I*_*V*_ and *I*_*D*_ by *D* and *V*, respectively.

**Figure 2:**
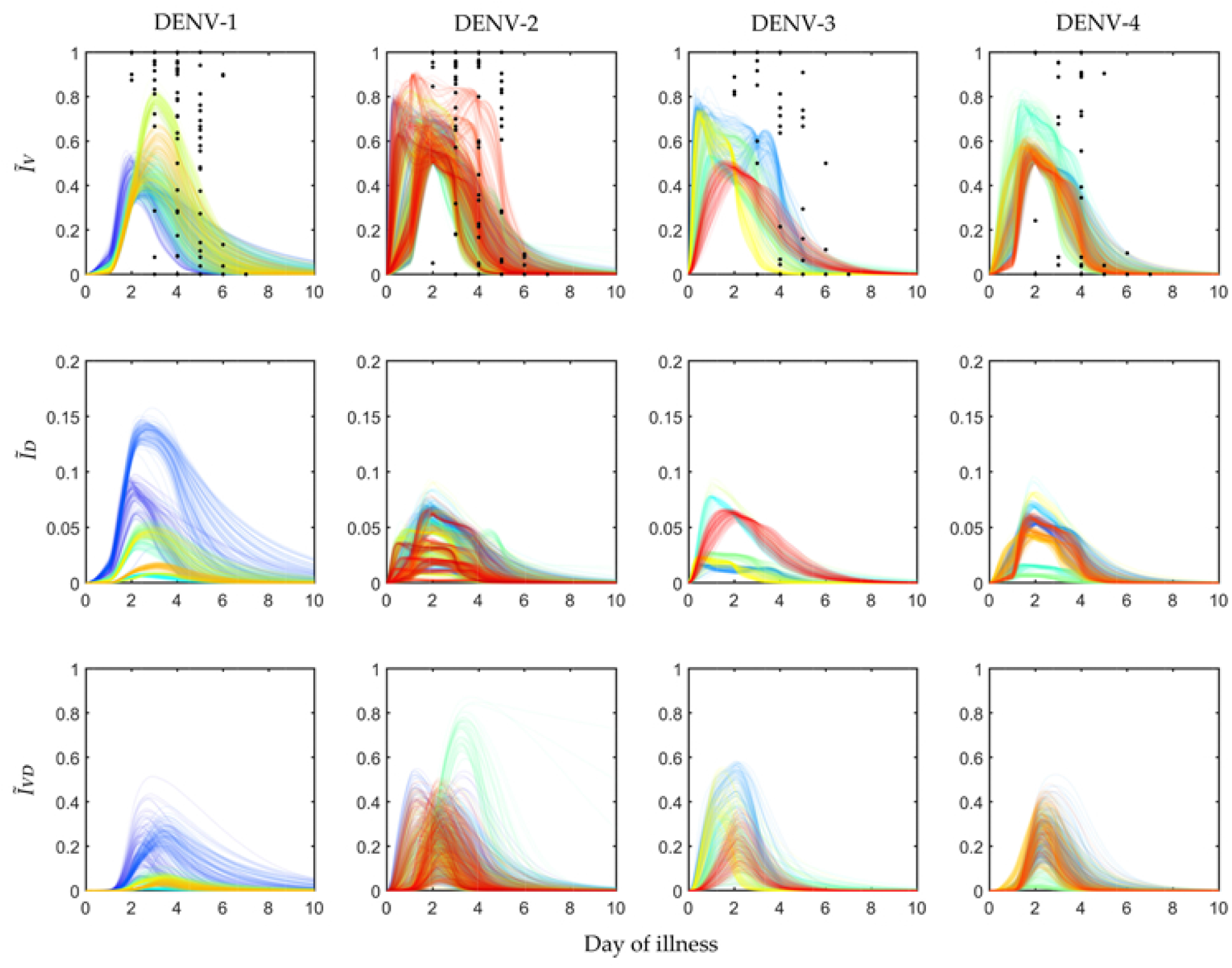
Population of models for infected mosquitoes: The dengue virus infected mosquito data for 408 exposure experiments with 208 hospitalised dengue patients reported in Nguyen *et al* ^32^ are calibrated to construct serotype-specific population of models (POMs). The biomarker data for available viral load in mosquito abdomens are shown as black dots and the calibrated models are shown as coloured lines. The different coloured lines represent different patient models involved in the mosquito exposure experiments and lines in the same colour demonstrate the population of models for each patient model.

In Fig 3, the selected parameters for the calibrated models are shown in box plots for the four dengue serotypes. During the calibration of POMs with the reported biomarker data, the different rate constants of mosquito reproduction (*A*), infection by host viraemia (*ϕ*) and natural mosquito death (*μ*) have been sampled using Latin Hypercube Sampling and selected within the POMs according to Algorithm 1. The accepted model parameters are normalised by their values and included in the box plot as scattered distributions.

**Figure 3:**
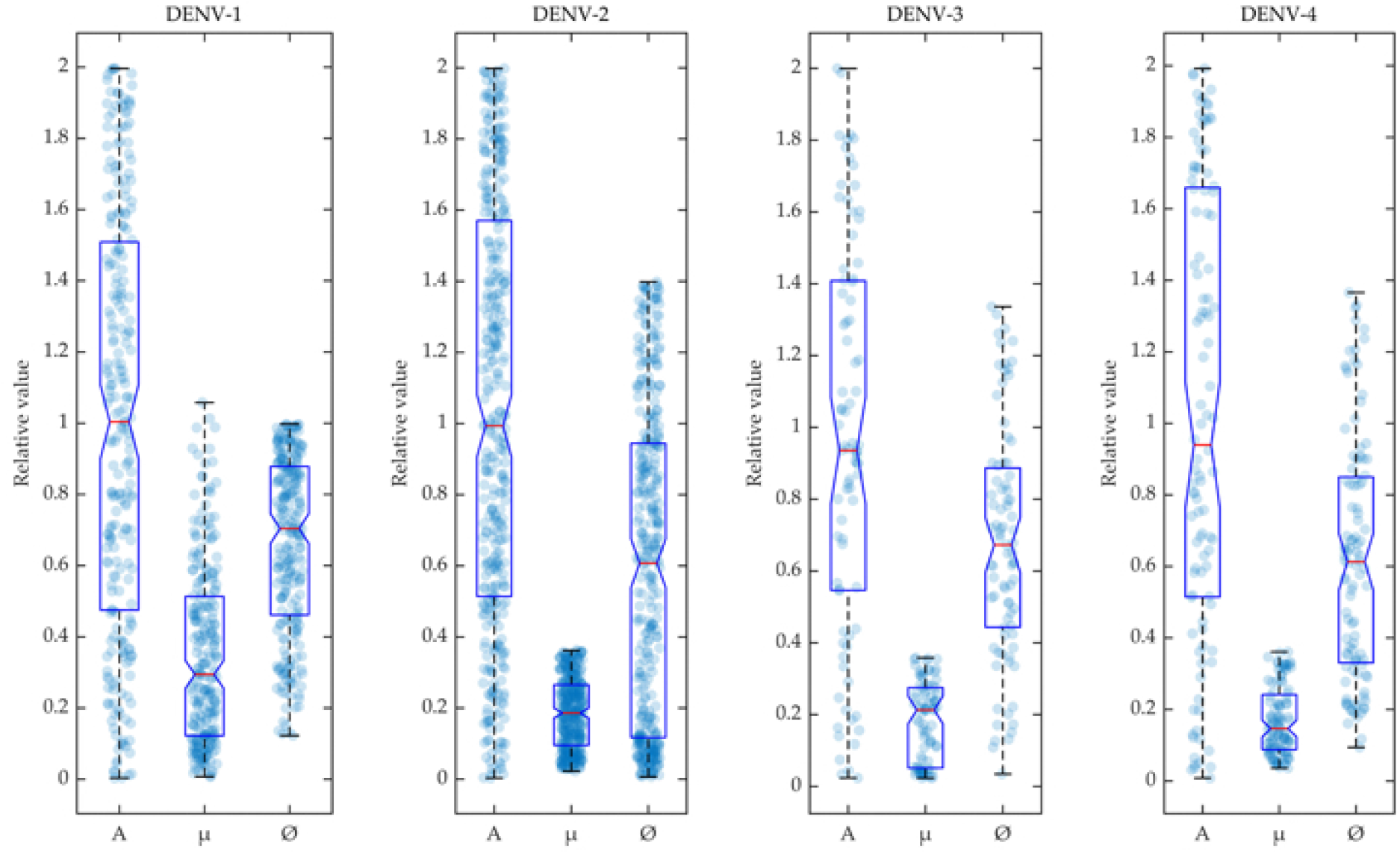
Variability in accepted model parameters: The parameters accepted in the POMs are shown in box plots for the four serotypes. The density of the parameters are shown in the scatter plots in the background of the box plots.

The successful human-to-mosquito transmission of DENV has been estimated by RT-PCR (reverse transcriptase polymerase chain reaction) of the viral titre from the mosquito abdomens ^32^. To correlate the patients’ plasma viraemia kinetics with the mosquito infection pattern, the blood-fed mosquitoes are harvested 12 days after the day of exposure. The efficiency of patient viraemia to infect the mosquito pool can be measured by the mosquito infectious dose (*MID*) in a quantitative way. The *MID* is a direct evaluation of the force of host-to-vector virus transmission. In the present study the plausible ranges of *MID*s are estimated from the four serotype-specific calibrated POMs (Fig 4). The infected mosquito fraction (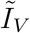) is plotted with respect to the Log10 of employed viral load (*V*(*t*)) from the within-host model ^31^. The distribution of the 50% mosquito infectious dose (*MID*_50_) is estimated for each serotype from the fraction of 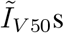 and the corresponding *V*(*t*) levels. As a patient with high viraemia is maximally efficient for transmission, very few transmissions are observed after day 5 of illness.

**Figure 4:**
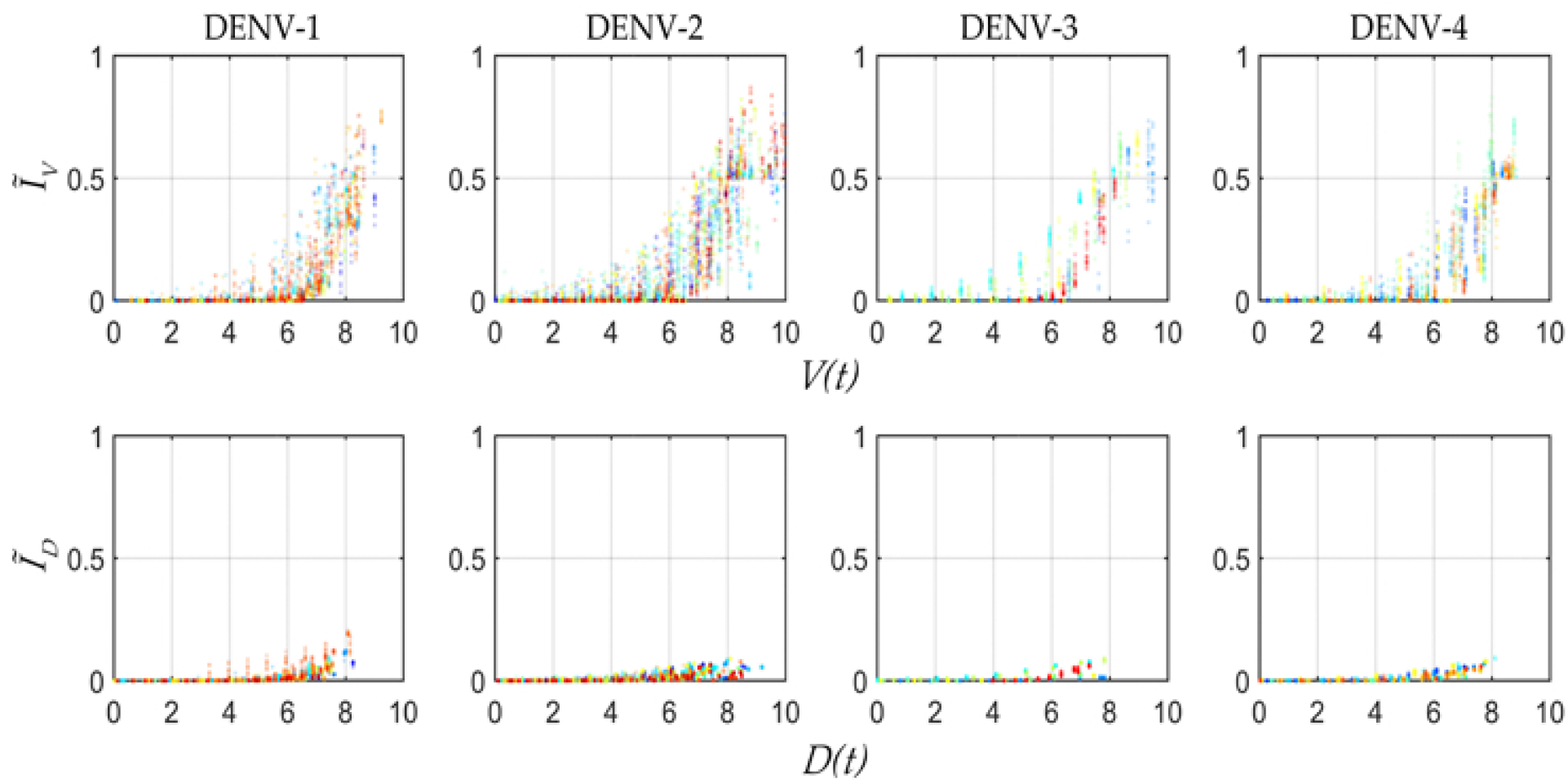
Mosquito infectious dose before applying control: The virus infected mosquitoes (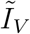) and Log_10_ values of the corresponding patient plasma viraemia (*V*(*t*)) are shown in scatter plots (colours represent different patients) to elucidate the ranges of the mosquito infectious dose. The infection of DI particles are also in terms of 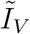 vs Log_10_ values of the corresponding patient DI particles profiles (*D*(*t*)). The range of MIDs are estimated from these scatter plots.

In Fig 5 the POMs for the four serotypes (across the columns) simulated with the controlled viraemia from the within-host model as the input are presented. If we observe the infection in mosquitoes by the virus (Fig 5), the 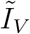 levels for all DENV serotypes are reduced more than 10-fold. For DENV-1 and DENV-2, the overall peaks of 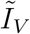 is close to 0.05, whereas the same peaks for the uncontrolled case (Fig 2) are near 0.9 and 0.85. We observe more efficient reduction in 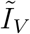 for DENV-3 and DENV-4. The controlled 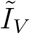 for DENV-3 is near 0.01 and for DENV-4 it is near The transmission of DI infection (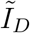) is increased significantly for all the serotypes, but it cannot dominate over the transmission of 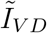. The most interesting results are observed in case of DENV-2 and DENV-3, where a majority of the patients appear with secondary infection. The utilization of the controlled viraemia does not greatly affect co-infection (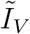) profiles during the virus infection reduction in those case of secondary infections. Although the co-infection population dynamics shows changes with respect to the population before intervention, the individual profiles in the whole population do not get affected. Notable reduction is observed in the case of DENV-1 and DENV-4 while DENV-2 and DENV-3 are still in a good state.

Fig 6 shows the controlled ranges of mosquito infectious dose (*MID*) for the four DENV serotypes. In most of the cases of DENV-1 and DENV-2, the 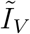 reduces ≈ 100−fold with the reduction in viraemia. However, some controlled *MID* points are observed as outliers in the population of DENV-2. If we observe these outlying points carefully, we can make a note that these points appear at the controlled peak values of the viraemias along the x-axis. The corresponding controlled 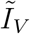 fractions (in cyan, Fig 5) have similar short-lived sharp peaks between day 0 and day 1 of illness. On this point we want to comment on the dengue transmission by the asymptomatic patients. For a particular value of viraemia, symptom-free dengue patients are significantly more infectious than clinically symptomatic patients ^47^. However, the outlying points in our controlled 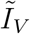 fractions are not in the region of significant transmission. On the other hand, with the same reduction in viraemia for DENV-3 and DENV-4, 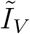 is reduced much more efficiently and they do not possess any such outlying points for 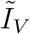 fractions (Fig 6).

**Figure 5:**
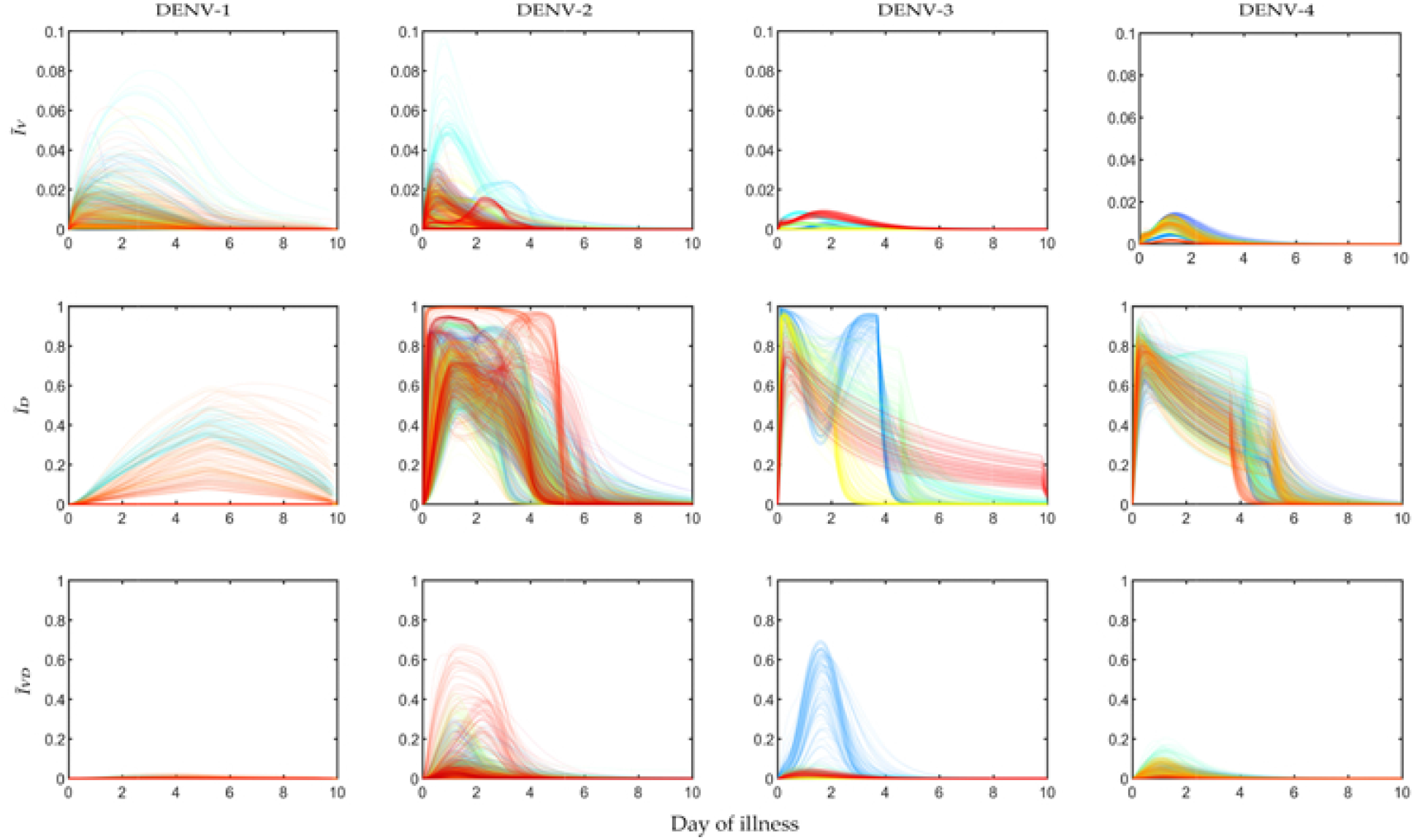
Controlled population of models for infected mosquitoes: Each serotype-specific POMs (In Figure 2) are considered with the controlled viraemia to contruct the controlled POMs (cPOMs) presented as: infected by virus only (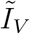), infected by DI only (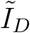) and infected by both (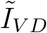), simulated with the controlled within-host plasma viraemia and DI particles in the populations for four dengue serotypes.

**Figure 6:**
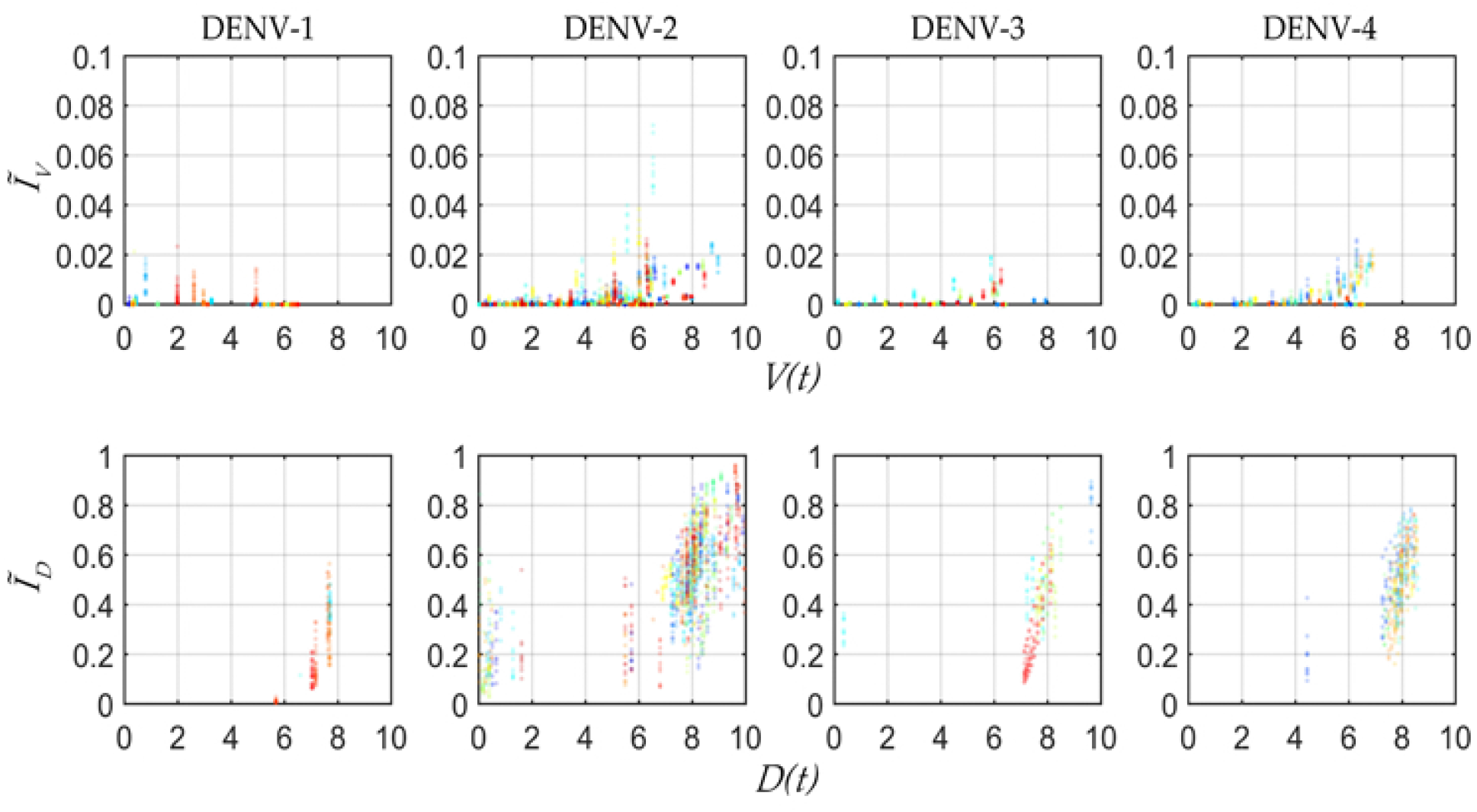
Controlled MIDs. Scatter plots show the attenuated MIDs and the transmission of DI particles with respect to the *Log*_10_ values of the plasma viraemia and DI particles after applying the control.

To estimate the efficiency of the DI particles-mediated intervention over virus transmission, we use a Jensen-Shannon divergence (JSD, 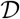), which is derived from the Jensen’s inequality and Shannon entropy ^48^. The JSD measure is a symmetric overall difference between two distributions. We calculate the JSD between the uncontrolled and controlled 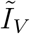 populations for each patient model on each day of illness as

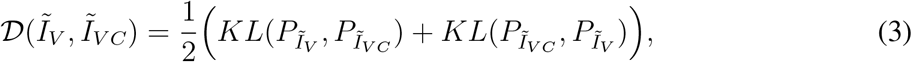

where, *KL* is the Kullback-Leibler divergence measure ^49^ between the probability distributions of uncontrolled and controlled infected mosquito fractions for each individual patient model. Here we denote the controlled 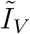 as 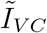. In Fig 7 the box plots in the inset produce the distribution of all the patients’ performance on viraemia reduction in terms of JSD. A greater JSD score implies more effective control and with that notion, DENV-2 and DENV-3 show more effectiveness with higher JSD values. Moreover, we know that an intervention strategy using excess DI particles will be called efficiently cost-effective if it can reduce maximum transmission with a minimum DI particles addition for minimum duration. Hence, we present the distribution of JSD with respect to the normalised control expense (A) and find DENV-2 with the most efficient control strategy for the present dataset ^31^.

## Discussion

The main goal of this paper is to calibrate a model for the infectedness of the patients to mosquito and to predict the effect of intervention with excess DI particles from the population of patients’ model to the mosquito transmission model. We use the method of population of models (POMs) with the viral load computed before and after applying bang-bang control in our previous model ^31^. As the reduction in virus transmission is directly related to the efficiency of the assisted transmission of DI particles, the same intervention strategy within the infected patients’ population may prevent the outbreak of dengue.

**Figure 7:**
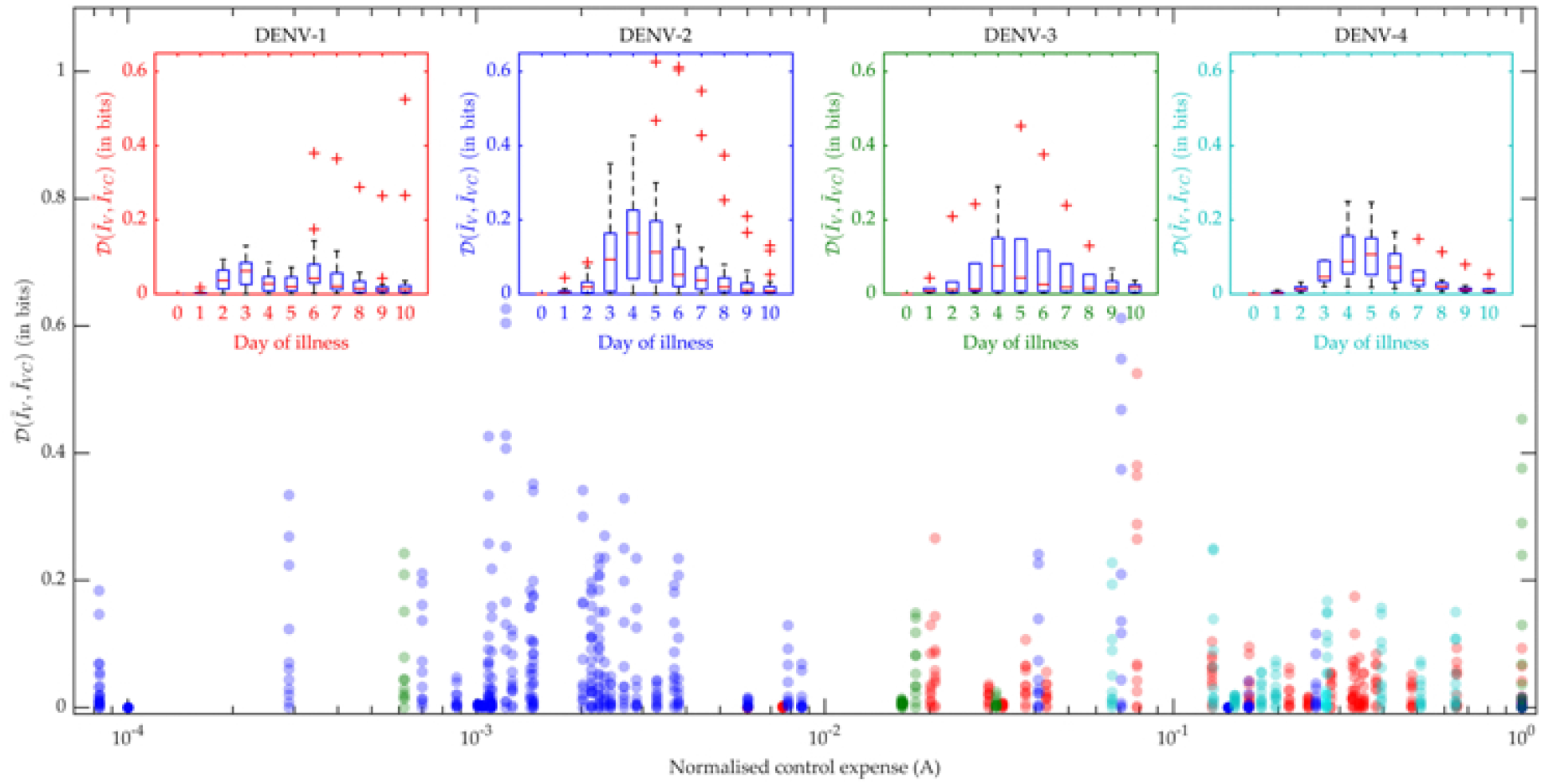
Control efficiency. The reduction in the human-to-mosquito virus transmission is evaluated using Jensen-Shannon divergence (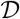) calculated between the distributions of the virus infected mosquitoes before (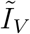) and after (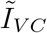) applying control. The box plots for each serotype explains the variation in the J-S divergence on each day of the febrile period. The scatter plot compares the efficiency of the applied control in reduction of the J-S divergence (red: DENV-1, blue: DENV-2, green: DENV-3, cyan: DENV-4) in terms of control expense (A).

It is clear from Fig 2 that as long as the host viraemia level stays very high, i.e., within 2-4 days of illness, the probability of transmission into mosquitoes is high. One of the reasons behind this phenomenon is the requirement of high viral load (see Fig 4) in the host blood for successful systematic replication through the tissue barriers in the mosquito body. Another reason is the shorter mosquito life cycle (approx. 15 days) that terminates mostly before the mosquito becomes infected. As Fig 4 cannot interpret the relative timescales of the host plasma viraemia and mosquito infection, we have to compare the time profiles of the mosquito infections (see Fig 2) and viraemia profiles in ^31^. The rise of the viraemia-infectivity relation can be observed in the representative lines (purple line) in Fig 4. The dengue fever starts with a very high viral burden (about 10^5^ to 10^7^) on day 0 of illness and grows sharply during the first few days of the febrile period. During these days the transmission probability is high and the MID value is higher than the *MID*_50_. However, the virus assisted transmission of DI particles (*I_V D_*) persists for longer, even after the virus is mostly cleared, while the only DI particles transmission is much lower than the standard virus and the assisted transmission. We wish to explain the dynamics in terms of the replication competition in the within-host and within-vector dynamics. In the very early days of illness, the host plasma viraemia is mostly populated by the standard virus and they replicate and transmit through mosquito bites. During these days, the population of virus-infected host cells grows faster and releases virus. The DI particles and cells infected by both the standard and DI particles appear to grow with a delay^31^. That delay is also reflected here in terms of transmission. The *I*_*V D*_ mosquito pool starts to reach high values as soon as the *I*_*V*_ starts to fall.

As we have discussed, the primary and secondary infections in the within-patient models can be classified with the help of their viraemia time profiles. The same situation is also reflected here in the mosquito model. As the majority of the DENV-2 and DENV-3 infected patients were diagnosed with a secondary infection, the POMs for DENV-2 and DENV-3 in the current model show sharper growth in transmission than DENV-1 and DENV-4. In contrast, the *I*_*V D*_ transmission for DENV-2 and DENV-3 are lower than that in the case of DENV-1 and DENV-4. Fig 2 shows the density distribution of the *I*_*V D*_ trajectories for different serotypes. These results suggest that the more secondary infections occur, the more the standard virus will be transmitted with respect to DI particles.

The controlled profiles of the infection compartments of the mosquito pool show log scale reduction from the uncontrolled profiles. The maximum reduction is observed in the profiles of *I*_*V*_ after applying the control, whereas the *I*_*V D*_ profiles are not reduced completely. These results explain the underlying trade-off between the transmission of virus and DI particles. Our goal is to completely block the virus transmission with a persistent good level of DI particles transmission. However, the passage of DI particles is not possible without the assistance of the transmission of a helper virus. Hence, a high-efficiency passage of DI particles needs to allow a lower minimum level of virus to be transmitted. Nevertheless, the allowed level of viral transmission is not enough to produce dengue fever or endemic outbreak.

Although quantitative detection of the existence of viral load in different mosquito body parts is a common experiment, it is hard to detect if there exists any DI RNA. To tackle such problems, a predictive model may address a number of questions. We believe, this model is the first to explore the possible scenario of a mosquito pool, infected by blood-feeding themselves from dengue infected human host. We hope to develop another model on virus replication and the mechanism of natural occurrence of DI particles via genome deletion and mutation. This model will consider the multi-class queue of positive stranded RNA (full length and defective) and investigate the delay in terms of dengue control strategy.

## Acknowledgements

We thank Nguyen et. al. for providing the data collected at the Hospital for Tropical Diseases, HCMC, Vietnam. We thank all those engaged in the DARPA INTERCEPT Dengue TIPs for suggestions and discussions. We thank the High Performance Computing (HPC) and Research Support of QUT for the computational facility.

